# HiLDA: a statistical approach to investigate differences in mutational signatures

**DOI:** 10.1101/577452

**Authors:** Zhi Yang, Priyatama Pandey, Darryl Shibata, David V. Conti, Paul Marjoram, Kimberly D. Siegmund

**Affiliations:** Department of Preventive Medicine, Keck School of Medicine of the University of Southern California, Los Angeles, CA USA; Department of Pathology, Keck School of Medicine of the University of Southern California, Los Angeles, CA USA

**Author notes:** Corresponding author: Zhi Yang.

## Abstract

We propose a hierarchical latent Dirichlet allocation model (HiLDA) for characterizing somatic mutation data in cancer. The method allows us to infer mutational patterns and their relative frequencies in a set of tumor mutational catalogs and to compare the estimated frequencies between tumor sets. We apply our method to somatic mutations in colon cancer with mutations classified by the time of occurrence, before or after tumor initiation. Applying the methods to 16 colon cancers, we found significant associations between the relative frequencies of mutational patterns and the time of occurrence of mutations. Our novel method provides higher statistical power for detecting differences in mutational signatures.

## INTRODUCTION

A variety of mutational processes occur over the lifetime of an individual, and thereby uniquely contribute to the catalog of somatic mutations observed in a tumor. Some processes leave a molecular signature: a specific base substitution occurring within a particular pattern of neighboring bases. A variety of methods exist to discover mutational signatures from the catalog of all somatic mutations in a set of tumors, estimating the latent mutational signatures as well as the latent *exposures* (i.e., fraction of mutations) each signature contributes to the total catalog. The first large study of mutational signatures in cancer identified variation in mutational signatures and mutational exposures across 21 different cancer types (Alexandrov et al., 2013). To better understand the sources of variation in the mutational exposures across cancers, our interest is in statistical methods used to characterize these latent mutational exposures across different cancer subtypes. Moreover, by classifying mutations by their time of occurrence, before or after tumor initiation, we can investigate whether new mutational processes occur during tumor growth.

Previous studies interested in comparing mutational exposure estimates between different groups of tumor catalogs conducted a post hoc analysis. The analysis proceeded in two stages. First, they performed one of the several different approaches for mathematically extracting the latent mutational signatures and their exposures from the mutational catalogs (see Baez-Ortega and Gori (2017) for a review of such methods). Later, they conducted an independent test of association between the point estimates of the mutational exposures and external covariates. Examples of 1covariates included cancer subtype, or patient history of alcohol or tobacco use. A common choice for the second stage test is a Wilcoxon rank-sum test (Mann and Whitney, 1947; Network et al., 2017; Chang et al., 2017; Hillman et al., 2017; Letouzé et al., 2017; Meier et al., 2018; Haradhvala et al., 2018; Qin et al., 2018; Olivier et al., 2019; Guo et al., 2018). However, the variation of the exposure estimates is affected by two factors, the number of mutations in the tumor and the variation in exposure frequency in the patient population. The former, the number of mutations in the tumor, affects the accuracy of the exposure estimates. The application of the Wilcoxon rank-sum test on the exposure estimates does not take into consideration their accuracy, which can lead to loss of efficiency and test power. We address this by introducing a unified parametric model for testing variation of mutational exposures between groups of mutational catalogs, where the exposure frequencies are modeled using a Dirichlet distribution.

We propose a hierarchical latent Dirichlet allocation model (HiLDA) that adds an additional level to the latent Dirichlet allocation (LDA) model from Shiraishi et al. (2015). Shiraishi’s model, like the majority of deconvolution approaches, focuses on signatures for single-nucleotide substitutions, characterizing the mutation types by context, using local features in the genome such as the pattern of flanking bases and possibly the transcription strand. For both model parsimony and interpretation, we choose to extend their LDA model. First, it requires fewer parameters than competing methods, giving it higher power to detect patterns 5 bases in length compared to other models that consider only 3-base contexts (Shiraishi et al., 2015). Second, signature visualization methods lead to easy interpretation; an example is the common C>T substitution at CpG sites instead of the more complicated NpCpG patterns that appear when using the trinucleotide context. Like the LDA model, HiLDA retains all the functionality for estimating both the latent signatures and the latent mutational exposure of each signature for each tumor catalog. Our newly-added hierarchical level allows HiLDA to simultaneously test whether those mean exposures differ between different groups of catalogs while accounting for the uncertainty in the exposure estimates. Additionally, we can now parse out differences in group means in the presence of differences in group variances, which is not tenable when using post hoc nonparametric location-scale tests.

In this paper, we use HiLDA to study the association between the mutational exposures and the time of mutation occurrence in tumorigenesis. We classify cancer mutations into *trunk* or *branch* mutations: trunk mutations being those that occur before growth of the tumor, while branch mutations are those that occur during the tumor expansion process. A test of whether mutational exposures differ by time of mutation occurrence will allow us to assess whether new mutational processes occur following the transformation of the first cancer cell.

## METHODS

### Hierarchical Bayesian Mixture Model

We introduce a hierarchical latent Dirichlet allocation model (HiLDA) using the following notation, also summarized in Table 1. Let *i* index the mutational catalog and *j* the mutation. The nucleotide substitutions are reduced to six possible types (C>A, C>T, C>G, T>A, T>C, T>G) to eliminate redundancy introduced by the complementary strands. Each observed mutation is characterized by a vector, ***X***_*i,j*_ describing the nucleotide substitution (e.g. C>T) and a set of genomic features in the neighborhood. Example features include the base(s) 3*′* and 5*′*of the nucleotide substitution (C, G, A, T), and the transcription strand (+,-). Each observed feature characteristic, *x*_*i, j,l*_ for mutation feature *l*, takes values in the set {1, 2, …, *M*_*l*_*}* (where *M*_*l*_ = 6 for the nucleotide substitution, or 4 for a flanking base, and 2 for the transcription strand).

**Table 1.** List of notation.

| Notation | Description |
| --- | --- |
| $I$ | Total number of mutational catalogs (indexed by $i$) |
| $J_i$ | Number of observed mutations in $i$th mutational catalog (indexed by $j$) |
| $L$ | Number of features to include. Here, we use the nucleotide substitution, flanking bases and transcription strand (indexed by $l$) |
| $\mathbf{M}$ | Vector of the maximum numbers of possible values, $(M_1, \dots, M_L)$, for each mutation feature, (indexed by $M_l$), $M_1 = 6$ for nucleotide substitution, $M_2 = 4$ for flanking base, (A, C, G, T), $M_L = 2$ for transcription strand, $(+, -)$ |
| $K$ | Total number of mutational signatures (indexed by $k$) |
| $\mathbf{X}_{i,j}$ | Observed mutation characteristic vector, $(x_{i,j,1}, \dots, x_{i,j,L})$, for the $j$th mutation from the $i$th mutational catalog (indexed by $x_{i,j,l}$) |
| $z_{i,j}$ | Index of the latent assignment for $\mathbf{X}_{i,j}$, $z_{i,j} \in \{1, \dots, K\}$ |
| $\mathbf{q}_{i,k}$ | Probability vector of signature $k$ exposure in mutational catalog $i$, $(q_{i,1}, \dots, q_{i,K})$, with $\sum_k q_{i,k} = 1$ |
| $\mathbf{f}_{k,l}$ | Probability vector of observing any of $M_l$ elements for $l$th mutation feature, $\mathbf{f}_{k,l} = (f_{k,l,1}, \dots, f_{k,l,M_l})$ with $\sum_{m_l} f_{k,l,m_l} = 1$ |
| $\mathbf{F}_k$ | A tuple of probability vectors with length $L$, $(\mathbf{f}_{k,1}, \dots, \mathbf{f}_{k,L})$ |
| $\mathbf{g}$ | A vector indicating group membership of the samples. ($g_i \in \{1, 2\}$ for each sample $i$) |
| $\boldsymbol{\alpha}$ | A tuple of concentration parameters of a Dirichlet distribution with length $K$, $(\alpha_1, \dots, \alpha_K)$, where the dispersion $\phi = \sum_k \alpha_k$ |
| $\boldsymbol{\mu}$ | A tuple of expected values of $\mathbf{q}$ of a Dirichlet distribution with length $K$, $(\mu_1, \dots, \mu_K)$, where $\sum_k \mu_k = 1$. |

We assume each mutation belongs to one of *K* distinct signatures. A specific mutational signature *k* is defined by an *l*-tuple of probability vectors, ***F***_*k*_, denoting the relative frequencies of the *M*_*l*_ discrete values for the *l* features, i.e., a vector ***f*** _*k,l*_ for the *M*_*l*_ values corresponding to feature *l*. We let *z*_*i, j*_ denote the unique latent assignment of mutation ***X***_*i, j*_ to a particular signature. Then, given the signature to which a mutation belongs, the probability of observing a mutational pattern is calculated as the product of the mutation feature probabilities for that signature. Thus, for signature *k* we write *Pr*(***X***_*i,j*_|*z*_*i, j*_) = ∏_*l*_*f*_*k,l*_ (*x*_*i, j,l*_| *z*_*i, j*_). This assumes independent contributions of each feature to the signature. To model each multinomial distribution of ***f*** _*k,l*_, we use a non-informative Dirichlet prior distribution with all concentration parameters equal to one.

The unique personal exposure history of each individual leads to them having a particular (latent) vector, ***q***_*i*_, indicating the resulting contribution of each of the *K* signatures to that individual’s mutational catalog. These ***q***s are modeled using a Dirichlet distribution with concentration parameters ***α***, i.e., ***q***_*i*_ ∼*Dir*(***α***). Extending this model to the two-group setting, we allow the Dirichlet parameters to depend on group, 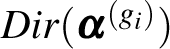, with *g*_*i*_ indexing the group corresponding to the *i*th catalog (*g*_*i*_ = 1 or 2). The mean mutational exposures, *E*(***q***_*i*_), denoted by 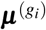, are represented by using the concentration parameters, i.e., 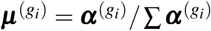.

With this extension, we can infer differences in mutational processes between groups of catalogs by testing whether the mean mutational exposures differ between the two sets, i.e., at least one 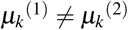. The likelihood and prior of the multi-level model is specified as follows,

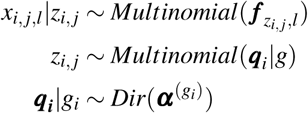

For full details see See Text S1. and Fig. S2..

### Testing for Differences in Signature Exposures

To characterize the signature contributions for different sets of tumor catalogs, we wish to conduct a hypothesis test that there is no difference in mean exposures versus the alternative that the mean exposure of at least one signature differs between the two groups, i.e. *H*_0_ : ***µ***^(1)^ = ***µ***^(2)^ vs. *H*_1_: at least one 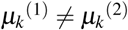. We propose both local and global tests, implemented in a Bayesian framework. The former provides signature-level evaluations to determine where the differences in mean mutational exposures occur, while the latter provides an overall conclusion about any difference in mean mutational exposures. The details of our implementation are given in our Just Another Gibbs Sampler (JAGS) scripts and Source code is freely available in Github at https://github.com/USCbiostats/HiLDA (Plummer et al., 2003).

### A local test to identify signatures with different exposures

We propose a signature-level (local) hypothesis test to allow us to infer which signature(s) contribute a different mean exposure to the mutational catalogs across tumor sets, i.e., 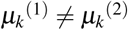. To measure the difference between mean signature exposure vectors, we implement HiLDA by specifying two Dirichlet distributions, *Dir*(***α***^(1)^) and *Dir*(***α***^(2)^), as priors for the distribution of mutational exposures ***q***_*i*_ of each group (Spiegelhalter et al., 2003). Using this formulation, the difference between the two groups of the mean exposure of signature *k* is calculated as,

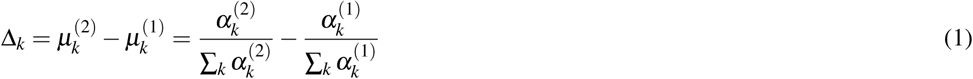

For all parameters,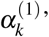 s and 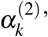 s, we use independent, non-informative gamma distribution priors with a rate of 0.001 and shape of 0.001; this results in a mean of 1 and variance of 1000. So,

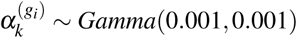

We estimate parameters via Markov chain Monte Carlo (MCMC) using two chains (Carlin and Chib, 1995). We assess convergence of the two MCMC chains using the potential scale reduction factor (Rhat) in Gelman et al. (1992), which is required to be less than or equal to 1.05 for all parameters in order to conclude that the MCMC run has converged. After obtaining the posterior distribution of the differences (i.e., of Δ_*k*_), there are two possible approaches to performing inference. We can: 1) use the Wald test to compute the *P*-value using the means and standard errors of the posterior distribution for Δ_*k*_; 2) determine whether the 95% credible interval of the posterior distribution for Δ_*k*_ contains zero.

### A global test using the Bayes factor

We also propose a global test to provide an overall conclusion on whether the mean exposures differ between groups of catalogs. It uses the Bayes factor, the ratio of posterior to prior odds in favor of the alternative (*H*_1_: at least one 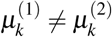, *k*=1, …, *K*) compared to the null (*H*_0_: ***µ***^(1)^ = ***µ***^(2)^), to indicate the strength of evidence that they do differ, without explicit details on how they differ. Thus, we can calculate the Bayes factor as:

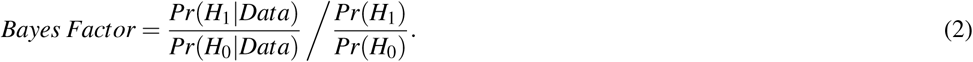

Since the likelihood is analytically intractable, the Bayes factor is calculated via MCMC (Carlin and Chib, 1995). In order to estimate the Bayes factor, during the MCMC analysis, a single binary hypothesis index variable is used to indicate which hypothesis explains the observed data (Lodewyckx et al., 2011). The parameters of two Dirichlet distributions, *Dir*(***α***^(1)^) and *Dir*(***α***^(2)^), are drawn from the same prior if the index takes the value 1, whereas they are drawn from different priors if it takes the value 2. Initially, the prior hypothesis odds is set to be 0.5/0.5 = 1, which means that both hypotheses are assumed equally likely under the prior. In order to improve computational efficiency in extreme situations in which one hypothesis dominates the other, we can use a different prior odds value (Carlin and Chib, 1995).

### Two-stage Inference Methods using the Point Estimates of mutational exposures

An alternative approach is to perform hypothesis testing using point estimates of the mutational exposures, 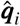, in a two-stage analysis, which we refer to as the “two-stage” method (TS). We used the R package **pmsignature** to estimate 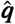 (Shiraishi et al., 2015). Other methods are also available, but we selected **pmsignature** for the purpose of comparisons to the results from HiLDA since it assumes the same model for estimating signatures under independence of features. We summarize the steps of the TS method as follows:

1. Jointly estimate the vectors of mutational signature exposures, ***q***_*i*_, for each mutational catalog.
2. Test for differential mutational exposures for signature *k* by performing the Wilcoxon rank-sum test on the 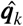.

However, we note that the Wilcoxon rank-sum test in stage 2 is also sensitive to changes in variance across the two groups, which might lead to significant results even when there has been no change in mean exposures (Kasuya, 2001; Ruxton, 2006). We implemented the two-stage method using R version (R Core Team, 2017). A two-sided P value of less than 0.05 was considered statistically significant.

### Choosing the Number of Signatures

The number of signatures, *K*, needs to be determined prior to any of the above analyses. We adopted the method of Shiraishi et al. (2015) to determine *K*. Their method is based on the following criteria:

1. The optimal value of *K* is selected over a range of *K* values such that the likelihood remains relatively high while simultaneously having relatively low standard errors for the parameters.
2. Pairwise correlations between any two signatures (the *k*th signature and the *k*^*′*^th signature, say) are measured by calculating the Pearson correlation between their estimated exposures across all samples, (i.e., the correlation between 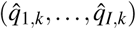 and 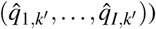). *K* is chosen such that no strong correlation (i.e., >0.6) exists between any pair.

For full details see Shiraishi et al. (2015).

### Application to Tumor Evolution

#### USC Colon Cancer Data

Our goal is to identify whether any new mutational signatures occur during colon cancer growth that distinguish cancer evolution from normal tissue evolution. To achieve this, we classify somatic mutations into two catalogs according to time of occurrence: those that accumulated between the time of the zygote and the first tumor cell, which we call trunk mutations, and those that occur *de novo* during tumor growth, which we refer to as branch mutations. We then estimate mutational signatures in the two sets of catalogs and test whether the mean mutational exposures differ between them.

We analyzed a total of 16 colon tumors. Tumor and adjacent normal tissue were subject to whole exome sequencing, and somatic mutations called using the GATK pipeline and MuTect (details below). Somatic mutations in the tumors were defined as nucleotide variants that were detected in tumor tissue but did not also appear in the patient-matched normal tissue. We used multi-region tumor sampling to allow us to distinguish between trunk from branch mutations (Siegmund and Shibata, 2016). Each tumor was sampled twice, with bulk tissue samples taken from opposite tumor halves. We classified somatic mutations appearing in both tumor halves as trunk, because only trunk mutations are likely to appear in both tumor halves, while mutations found on only one side of a tumor were labeled as branch. This approach has previously been shown to be 99% sensitive for calling trunk mutations and 85% sensitive for calling branch mutations (Siegmund and Shibata, 2016). Fifteen of the 16 tumors were previously analyzed in a study of cell motility (Ryser et al., 2018).

The sequence data were processed using the GATK pipeline version 3.7 (DePristo et al., 2011) and somatic mutations called with MuTect version 1.1.7 (Cibulskis et al., 2013), applying the quality filters KEEP (default parameters) and COVERED (read depth of 14 in tumor and 10 in matched normal – use of a lower coverage threshold in normal tissue is as recommended in (Cibulskis et al., 2013)). We excluded any mutations that either had an allele frequency less than 0.10, because sequencing errors are more common among low-frequency mutations (Cibulskis et al., 2013), or that were not also found by Strelka (Saunders et al., 2012), which we used as a confirmatory control. Somatic mutations on chromosomes 1 to 22 were used for mutational signature analysis.

## RESULTS

### Application to Tumor Evolution

A total of 12,554 somatic single-nucleotide substitutions were identified, with a median of 277 per sample (range: 82 – 1,762) (See Table S3.). One tumor with microsatelite instability has more than double the number of somatic mutations (1751 side A, 1762 side B) than any of the remaining 30 catalogs (all *<*750 mutations). In our first analysis, we compared the mutational exposures in side A to those in side B. If the tumors represent a single clonal expansion, we would expect similar mutational exposure frequencies in the two catalogs from the same tumor. Indeed, this is what we found (Table 2).

**Table 2.** Comparing mutational exposures from two sets of mutational catalogs, Side A and Side B, in the USC data.

|  | Side A - Side B | HiLDA-CI | HiLDA-Wald | TS-Wilcoxon |
| --- | --- | --- | --- | --- |
| Tests<sup>a</sup> | Coef. | [95% C.I.]<sup>b</sup> | p value | p value |
| $\Delta_1$ | 0.002 | [-0.079, 0.083] | 0.9863 | 0.7804 |
| $\Delta_2$ | 0.000 | [-0.029, 0.029] | 0.9875 | 0.8965 |
| $\Delta_3$ | -0.002 | [-0.083, 0.086] | 0.9608 | 0.9852 |
| $H_0 : \Delta_1 = \Delta_2 = \Delta_3 = 0$ | | Bayes Factor$_{M_2/M_1} = 0.021$ | | |

<sup>a</sup> $\Delta_k = \frac{\alpha_k^{(2)}}{\sum_k \alpha_k^{(2)}} - \frac{\alpha_k^{(1)}}{\sum_k \alpha_k^{(1)}}$ , the difference in the mean exposure of signature $k$ in group 1 and 2.
<sup>b</sup> 95% credible interval from the posterior distribution.

We identified a median of 174 trunk and 186 branch mutations per tumor. The numbers ranged from 49 to 1,578 trunk mutations and from 66 to 503 branch mutations (Fig. 1A). Interestingly, the microsatellite instable tumor had the most trunk mutations, but not the most branch mutations, suggesting that during tumor growth the mutation frequency is similar in microsatellite stable and instable tumors. Fig. 1B shows that the C>T substitution is most common in all trunk catalogs, and most branch catalogs. The spontaneous deamination of methylated Cs in CpGs is known to contribute to hotspots of C>T mutation in the genome.

**Figure 1.**
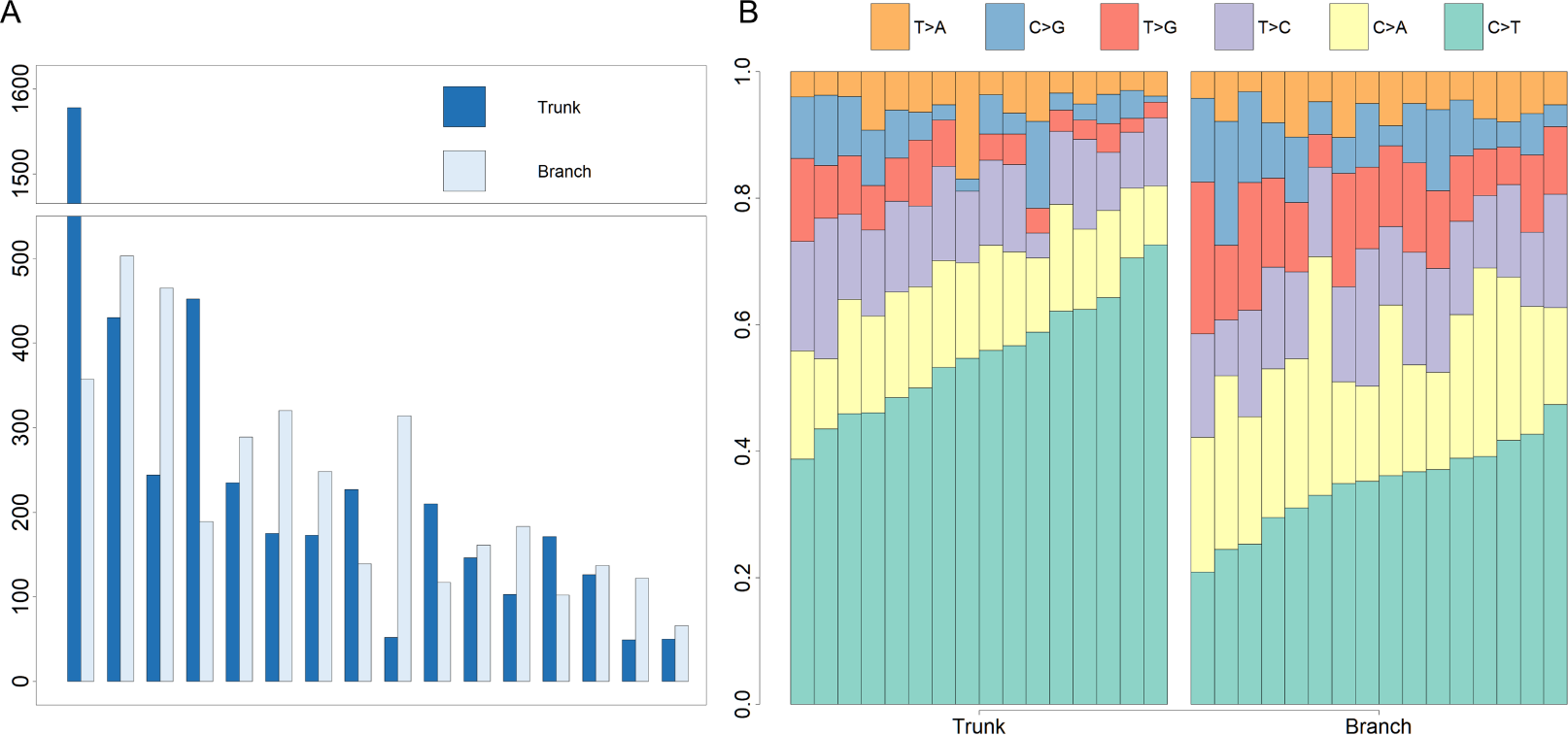
The numbers of somatic mutations in 32 mutational catalogs obtained from 16 colon cancer patients in the USC data and their mutation spectra. (A) The number of somatic mutations in 16 tumors, each of which contributes 2 mutational catalogs denoted as trunk (dark blue) and branch (light blue). (B) The percentage bar plot of relative frequencies for six substitution types in the 16 trunk mutational catalogs (left side) and the 16 branch mutational catalogs (right side).

We identified three mutational signatures in our data (see Fig. S4.). Those three signatures, and their corresponding exposures, are depicted in Fig. 2. The signature shown in the yellow box in the same figure, involving C>T mutations at NpCpG sites, resembles signature 7 in Shiraishi et al. (2015), where it was identified in 25 out of 30 cancer types and likely relates to the deamination of 5-methylcytosine (‘aging’); the signature in the orange box, involving T>G mutations at GpGpTpGpN sites, is novel; the third signature, in the red box, is qualitatively similar to signature 17 in Shiraishi et al. (2015), reflecting a signal specific to colorectal cancers. The pairwise cosine similarities between pairs of signatures are 0.12, 0.01, and 0.02 which are rather dissimilar from each other given the [0, 1] range for cosine similarity. Using HiLDA, we test whether the three signatures differ in mean exposure between trunk and branch mutations.

**Figure 2.**
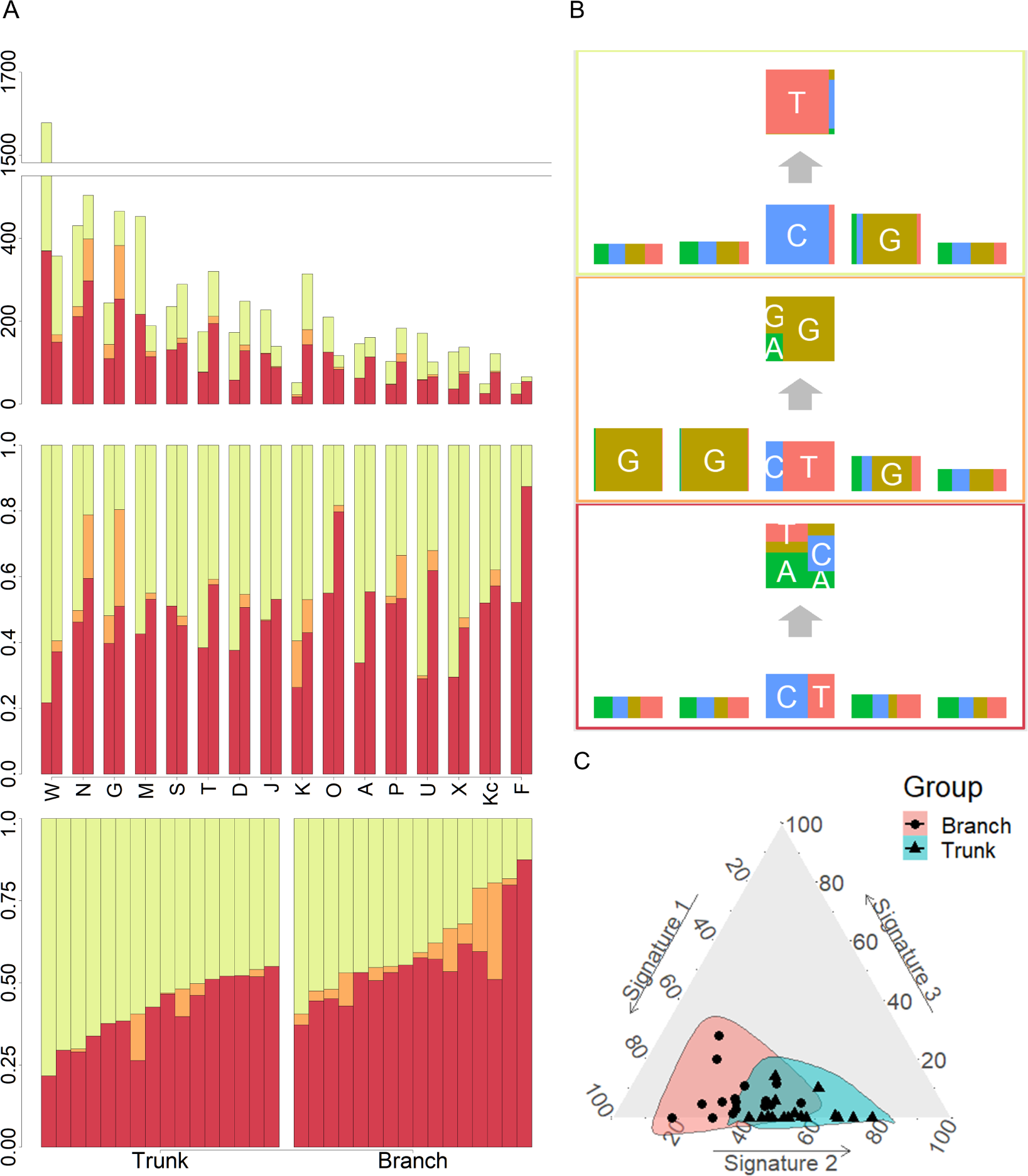
mutational exposures and three mutational signatures from the analysis of 16 trunk mutational catalogs and 16 branch mutational catalogs in the USC data (16 colon cancer patients). (A) From top to bottom, the three plots represent the somatic mutation counts, the corresponding mutational exposures, and the mutational exposures sorted by group (trunk/branch) and the exposure frequency of the first signature (yellow). (B) The three mutational signatures with four flanking bases. (C) The distributions of mutational exposures of the three mutational signatures highlighted by group, where the branch mutational catalogs are highlighted as pink and the trunk ones are highlighted as blue.

Our global test strongly suggests that, in our data, the signature exposures statistically differ between trunk and branch catalogs (Bayes Factor 1265.0). Each of the individual signatures (depicted in Fig. 2B) is found to differ in exposure between the two sample groups, a conclusion supported by both HiLDA and the two-stage method (Table 3). From Fig. 2A, it is evident that the exposures of the first (‘aging’) signature in trunk mutations is almost always greater than that for the matching catalog of branch mutations, which is intuitively consistent with the fact that trunk mutations may well reflect an accumulation of mutations over the life of the subject, whereas branch mutations are accumulated only after tumor initiation. For the previously unseen signature, the higher exposures in branch catalogs might suggest that this signature’s underlying mechanism for generating mutations might be associated with the processes occurring during tumor evolution as opposed to normal development. From Fig. 2C, we observed that the distributional ranges of the two groups of mutational exposures have some overlaps, but that the centers of each group, i.e., the means of mutational exposures, are clearly deviated from each other. However, the distributional radii, indicating the variances of mutational exposures, do not substantially differ between the groups.

**Table 3.**
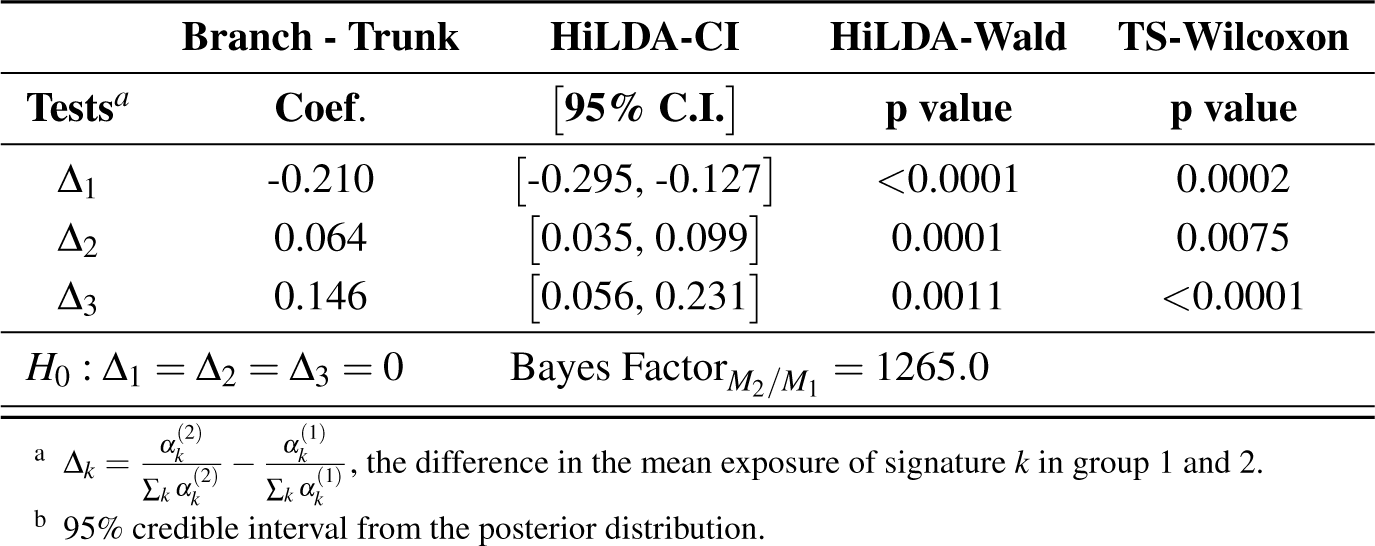
Comparing mutational exposures in colorectal cancer from two sets of mutational catalogs, trunk and branch, in the USC data.

We sought to validate the discovery of the previously unseen signature using both targeted sequencing data from the same tumor set (Siegmund and Shibata, 2016) and using publicly available data from the Cancer Genome Atlas. Four T>G substitutions that we assigned to the previously unseen signature were part of an independent validation set of mutations subjected to targeted, high-coverage Ampliseq technology (Siegmund and Shibata, 2016); all four of these T>G substitutions failed to validate. Further, a systematic analysis of data from the Cancer Genome Atlas Williams et al. (2016) also did not find evidence for this signature. Therefore, we cannot rule out that the signature is the result of sequencing error. We now go on to assess the reliability of results using a simulation study.

### Simulation Study

We conducted a simulation study to assess the performance of both HiLDA and the two-stage approach in terms of the false-positive rate (FPR) and true-positive rate (TPR), in local, univariate tests of the difference in mean exposure between two groups of mutational catalogs. In order to assess the functionality of the methods in a setting similar to that of the USC data, we simulate somatic mutations directly using the estimated signatures (***f***_*k*_) from Fig. 2 for the same number of mutational catalogs (two groups of 16 catalogs each) and somatic mutations per catalog (*J*_*i*_ in S3 Table). The mutational exposures (***q*_*i*_**) were indirectly used to derive the concentration parameters of the Dirichlet distributions. The scenarios are as follows:

1. The two groups of mutational catalogs are from separate Dirichlet distributions with parameters ***α***^(1)^ = (9.2, 0.2, 7.5) and ***α***^(2)^ = (4.2, 0.6, 7.3). Here, the ***α****s* corresponds to the maximum-likelihood estimated parameters from the three exposure distributions in the trunk and branch mutational catalogs. This gives mean exposures of ***µ***^(1)^ = (0.54, 0.01, 0.44) and ***µ***^(2)^ = (0.35, 0.05, 0.60) in trunk and branch catalogs, respectively, for the aging signature, new signature, and random signature.
2. The two groups of mutational catalogs are from the same Dirichlet distribution, *Dir*(4.2, 0.6, 7.3), (so here we use the concentration parameters estimated from the branch mutational catalogs).

For each tumor, mutational exposures ***q*_*i*_**, are drawn from the Dirichlet distribution. Each set of probabilities parameterize a multinomial distribution later used to probabilistically choose the underlying mutational signature for a mutation (See Fig. S5.). Then, every mutation feature in the mutational pattern of the mutation is simulated independently from a corresponding multinomial distribution of the chosen signature. To estimate the FPRs, 1000 sets of data were simulated for scenario 2, when there is no difference in the exposure distribution between two groups of mutational catalogs. The two-stage method is slightly conservative for 1st and 3rd signatures (resulting FPRs of 4.3%, 5.2%, and 4.3%) when testing at the 5% significant level (Table 4). In comparison, HiLDA showed better control of the FPR by using the 95% credible interval of the posterior distributions (4.8%, 5.0%, and 5.1%). The Wald test also showed control of the FPR, except in the case of the rare signature when it was noticeably lower (3.7%), presumably due to the asymmetric posterior distribution.

**Table 4.** The false positive rates (n = 1,000) and true positive rates (n = 200) of both the two-stage method and HiLDA when applied to the simulated data.

| | Methods | $\Delta_1$ | $\Delta_2$ | $\Delta_3$ |
| --- | --- | --- | --- | --- |
| **FPRs** | HILDA-CI<sup>a</sup> | 4.8 % | 5.0% | 5.1% |
|  | HILDA-Wald<sup>b</sup> | 5.1 % | 3.7% | 5.4% |
|  | TS-Wilcoxon | 4.3 % | 5.2% | 4.3% |
| **TPRs** | HILDA-CI | 99.5% | 85.5% | 91.5% |
|  | HILDA-Wald | 99.5% | 80.5% | 92.5% |
|  | TS-Wilcoxon | 99.0% | 77.5% | 88.0% |

<sup>a</sup> Percentage of 95% credible intervals that exclude zero.
<sup>b</sup> Percentage of $P$ -values $< 0.05$ after applying the Wald test to the posterior distribution.

We then moved to scenario 1, where we simulated 200 data sets with a difference in mean exposures between the two groups of catalogs. Here, the statistical powers of both HiLDA and the two-stage method are high when detecting the difference in exposures for the 1st and 3rd signatures (Table 4). In contrast, for the 2nd signature, which has the lowest mean mutational exposure, the TPRs of all methods are lower (77.5% – 85.5%). By using the 95% credible interval of posterior distributions, HiLDA is able to distinguish a difference more often than the two-stage method (99.5% vs. 99.0%, 85.5% vs. 77.5%, and 91.5% vs. 88.0%). At the same time, using the credible interval resulted in higher TPRs compared to performing a Wald test (85.5 % vs. 80.5% for the 2nd signature). In summary, across tests involving these three mutational signatures, HiLDA provides higher statistical power to the TS method with a tendency of better improvement for signatures with lower mutational exposures, i.e., the power difference between HiLDA and the TS method is the highest (8%) for signature 2 with the lowest mean mutational exposures. The improvements in the power to detect the mean exposure difference is presumably due to the fact that HiLDA accounts for the uncertainty in the estimated mutational exposures and provides better model fit of the posterior distributions. All data were simulated in R 3.5.0 using the hierarchical Bayesian mixture model described in the methods section. All replicates reached convergence with an Rhat value less than 1.05 for each of the scenarios shown in Tables 2-4.

## DISCUSSION

In this paper, we present a new hierarchical method, HiLDA, that allows the user to simultaneously extract mutational signatures and infer mutational exposures between two different groups of mutational catalogs, e.g., trunk and branch mutations in our example application. Our method is built on the approach of Shiraishi et al. (2015), in which mutational signatures are characterized under the assumption of independence, and it is the first to provide a unified way of testing whether mutational processes differ between groups (here, between early and late stages of tumor growth). As a result, our method allows us to appropriately control the false positive rates while providing higher power by accounting for the accuracy in the estimated mutational exposures.

In our analysis of the USC data, which consist of 32 mutational catalogs extracted from tumors from 16 CRC patients, our method detected three signatures and indicated a statistically significant difference in mean exposures between groups. Two of the three signatures resemble signatures 7 and 17 found by Shiraishi et al. (2015). But, in addition, we found a novel signature Shiraishi et al. (2015). Signature 7 appears significantly more often in trunk mutations, which is consistent with the fact that it has previously been related to aging and trunk mutations have a longer time over which to occur (conceivably over the lifetime of the patient) than do branch mutations (which occur only during tumor growth). The new signature, which occurred more often in low frequency branch mutations, is very similar to a sequencing artifact described by Alexandrov et al. (2018) (cosine similarity = 0.93). We note that, for the USC data, the conclusions obtained from HiLDA were qualitatively the same as those obtained from the TS method. This is likely due to the relatively large effect size here (i.e., the difference of mean exposures between the two groups, divided by the standard errors of same, also known as the signal-to-noise ratio). (Alexandrov et al., 2018).

In the simulation study, both HiLDA and the TS approach were applied to datasets consisting of 16 tumors simulated under two scenarios to test for between group differences in the mutational exposures of three signature. The results indicated that our unified approach has higher statistical power for detecting differences in exposures for these signatures while controlling the 5% false positive rate. We suspect that the improvement in statistical power is because our unified method explicitly allows for the uncertainty of inferred mutational exposures, while the two-stage method fails to do so since it incorporates only the point estimates of those exposures. In addition, HiLDA provides posterior distributions for each parameter, thereby allowing construction of 95% credible intervals for parameters, and their differences, for example. As expected, this fully parametric approach is then more powerful than nonparametric approaches, which we see particularly when testing for differences in the rarer signatures.

We also note that the two-stage approach can become problematic with regards to controlling the type I error rate in particular scenarios, e.g., when the variances of exposures differ widely between the two groups. In our simulation study, we aimed to emulate the USC data, meaning that the exposure variances were quite similar between groups. Consequently, the Wilcoxon rank-sum test, the second-stage of the TS approach, was able to maintain a type I error of 5%. However, we note that the Wilcoxon rank-sum test is sensitive to differences found in either location or scale parameters of the two distributions being tested, i.e., it is sensitive to changes in both the mean and the variance. Therefore, when the variances change between two groups, the Wilcoxon rank-sum test may indicate statistically significant differences in distributions even when the means have not changed, (i.e., due to the difference in shape parameters rather than a difference between location parameters). In contrast, HiLDA explicitly focuses on detecting differences in means, and is robust to effects such as changes in variance. Consequently, when applying the TS method, one should be wary of interpreting significant results as evidence of a “difference in means” when using the TS method (as seems to be common Qin et al. (2018); Meier et al. (2018); Network et al. (2017)). We note that scenarios in which the variance of the estimated exposures differs will be common if the numbers of mutations per tumor varies between the two groups (e.g. when comparing microsatellite instable vs. microsatellite stable colon tumors), leading to an inflated false-positive rate if results from the TS method are interpreted as being evidence of a difference in means. (See Fig. S6. for a specific example of this.) We intend to explore this issue further in a future paper. We also intend to more fully investigate the factors that drive the ability to detect significant difference between groups across a much wider variety of scenarios.

