## Supplemental Text 1 for "HiLDA: a statistical approach to investigate differences in mutational signatures"

### Text S1.

The likelihood can be constructed with an additional hierarchy based on a latent allocation or mixed membership model, where the likelihood of the observed mutations is conditioned on the unobserved signature of origin. As in the statistical approach described in the work of (Shiraishi et al., 2015), we model the features under the assumption of independence, such that the complete data likelihood is a product of multinomial distributions over each mutation feature $\boldsymbol{f}_{k,l}$, in which the indicator variable denoting a mutation's signature of origin, $I[z_{i,j} = k]$, is unobserved. In their previous work, (Shiraishi et al., 2015) assumed that mutational exposures $\boldsymbol{q}$ are drawn from the same Dirichlet distribution $Dir(\boldsymbol{\alpha}) = Dir(\alpha_1, \ldots, \alpha_K)$:

$$Pr(\boldsymbol{q}|\boldsymbol{\alpha}) = \prod_{i=1}^{I} \left( \frac{\Gamma(\sum_k \alpha_k)}{\prod_k \Gamma(\alpha_k)} \prod_{k=1}^{K} q_{i,k}^{\alpha_k - 1} \right)$$

$$Pr(\boldsymbol{X}, \boldsymbol{Z}|\boldsymbol{f}, \boldsymbol{q}) = \prod_{i=1}^{I} \prod_{j=1}^{J_i} \prod_{k=1}^{K} \left( \prod_{l=1}^{L} f_{k,l}(x_{i,j,l}|z_{i,j}) q_{i,k} \right)^{I[z_{i,j}=k]}$$

where the elements of $\alpha_k$ can be used to calculate the means for $q_{i,k}$, i.e., $\mu_k = \frac{\alpha_k}{\sum_1^K \alpha_k}$. The denominator $\sum_1^K \alpha_k$ is a measure of precision for the means, so that the larger this measure, the tighter the distribution of $q_{i,k}$ around the means.

In our case, there are two groups of mutational catalogs, referred to as group 1 and group 2, respectively. Therefore, we extend the above model by using two different Dirichlet distributions, $Dir(\boldsymbol{\alpha}^{(1)}) = Dir(\alpha_1^{(1)}, \ldots, \alpha_K^{(1)})$ and $Dir(\boldsymbol{\alpha}^{(2)}) = Dir(\alpha_1^{(2)}, \ldots, \alpha_K^{(2)})$, from which mutational exposure parameters are drawn, using the same set of $k$ signatures for each individual (See S2 Fig). Thus, we write,

$$Pr(\boldsymbol{q}|\boldsymbol{\alpha}^{(1)}, \boldsymbol{\alpha}^{(2)}, \boldsymbol{g}) = Pr(\boldsymbol{q}|\boldsymbol{\alpha}^{(1)})^{I[g_i=1]} \cdot Pr(\boldsymbol{q}|\boldsymbol{\alpha}^{(2)})^{I[g_i=2]}$$

$$Pr(\boldsymbol{X}, \boldsymbol{Z}|\boldsymbol{f}, \boldsymbol{q}) = \prod_{i=1}^{I} \prod_{j=1}^{J_i} \prod_{k=1}^{K} \left( \prod_{l=1}^{L} f_{k,l}(x_{i,j,l}|z_{i,j}) q_{i,k} \right)^{I[z_{i,j}=k]}$$
