## Supplementary figures and images for "HiLDA: a statistical approach to investigate differences in mutational signatures"

### Supplemental Figure 2

#### Figure S2.

The HiLDA diagram in plate notation.

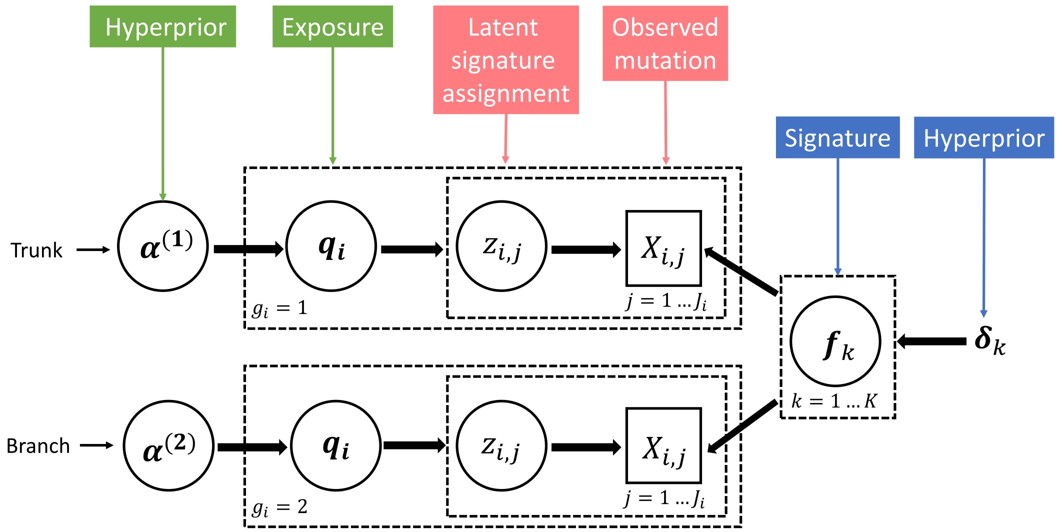
