## Supplemental Table 3 for "HiLDA: a statistical approach to investigate differences in mutational signatures"

### Table S3.

The number of somatic mutations in the 32 mutational catalogs from 16 colon tumors in the USC data, along with their associated trunk and branch mutations.

| Tumor | Side A | Side B | Trunk | Branch |
|---|---|---|---|---|
| W* | 1751 | 1762 | 1578 | 357 |
| N | 630 | 733 | 430 | 503 |
| M | 567 | 526 | 452 | 189 |
| G | 546 | 407 | 244 | 465 |
| S | 384 | 375 | 235 | 289 |
| T | 295 | 375 | 175 | 320 |
| D | 295 | 299 | 173 | 248 |
| J | 308 | 285 | 227 | 139 |
| O | 268 | 269 | 210 | 117 |
| A | 236 | 217 | 146 | 161 |
| U | 224 | 220 | 171 | 102 |
| K | 200 | 218 | 52 | 314 |
| P | 174 | 215 | 103 | 183 |
| X | 181 | 208 | 126 | 137 |
| Kc | 136 | 84 | 49 | 122 |
| F | 82 | 84 | 50 | 66 |
| Total | 6277 | 6277 | 4421 | 3712 |

* Sample W is microsatellite instable
