## Supplemental Figure 4 for "HiLDA: a statistical approach to investigate differences in mutational signatures"

### Figure S4.

The result of estimating mutational signatures for the USC data, including 16 trunk mutational catalogs and 16 branch mutational catalogs, using different values of $K$, the number of mutational signatures.

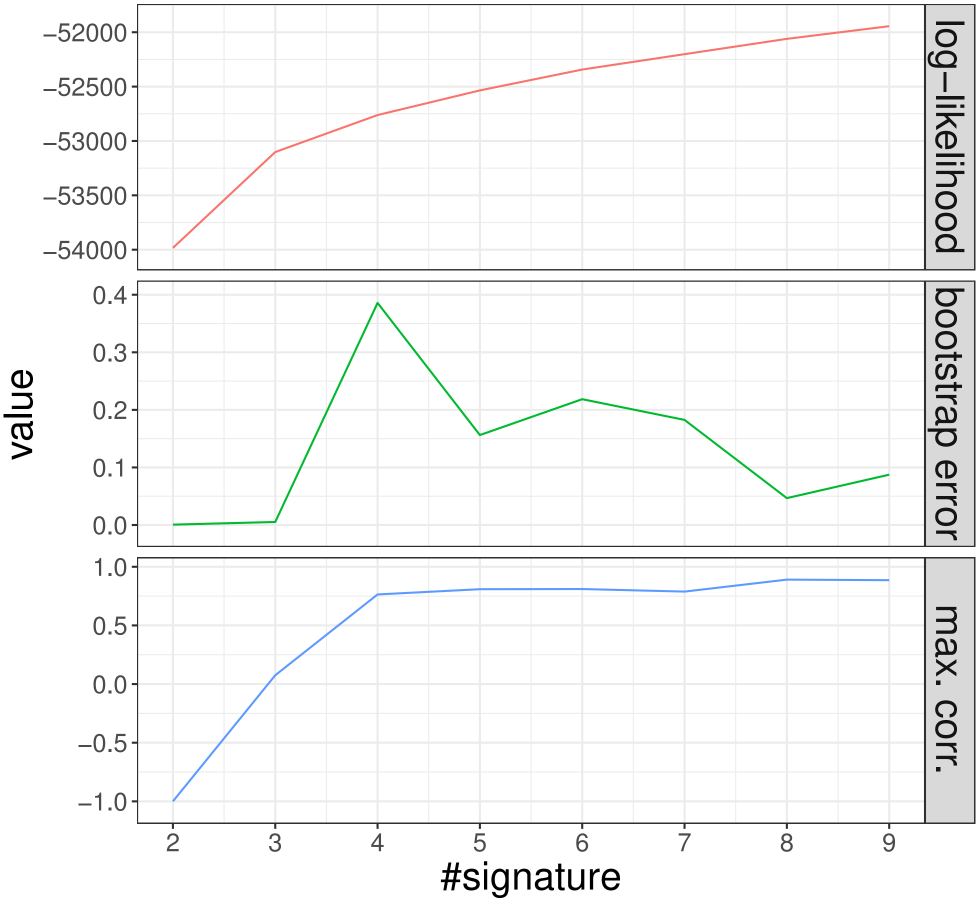


We fit our latent Dirichlet allocation model using the two bases flanking on both sides of our nucleotide substitution and select $K = 3$ signatures based on the three criteria proposed by (Shiraishi et al., 2015). We see that while increasing the number of signatures always results in an increase in the likelihood, the bootstrap-errors started to increase at $K = 4$ while the correlation between estimated signature fractions reaches high values at $K = 4$. This suggests that our 32 mutational catalogs can be explained using three signatures of mutational processes.
