## Supplemental Figure 5 for "HiLDA: a statistical approach to investigate differences in mutational signatures"

### Figure S5.

The generative process of a mutation in the simulation study is illustrated here. The black box outline is used to indicate the presumed generative process of a mutation, i.e., from the left to right, the simulated mutational exposures, the simulated signatures, and the simulated mutation features.

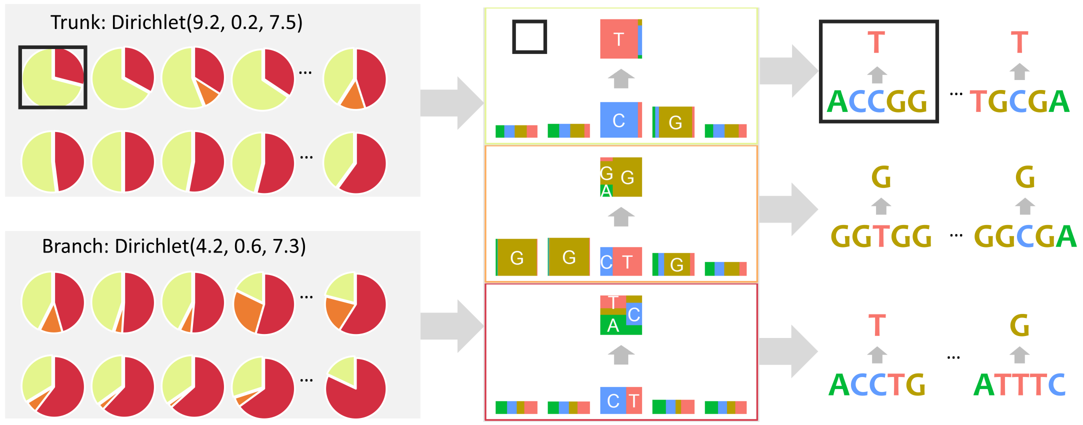


Let's take an example on how to simulate a somatic mutation in a trunk catalog. First, a multinomial distribution was drawn from the Dirichlet(9.2, 0.2, 7.5) for the mutational exposures of three signatures, which is represented by the pie chart. Then, the yellow signature was selected by randomly drawing from the previous multinomial distribution. Under the mutational patterns of this yellow signature, each mutation feature of this mutation was generated by using the multinomial distributions for the selected feature.
