## Supplemental Figure 6 for "HiLDA: a statistical approach to investigate differences in mutational signatures"

### Figure S6.

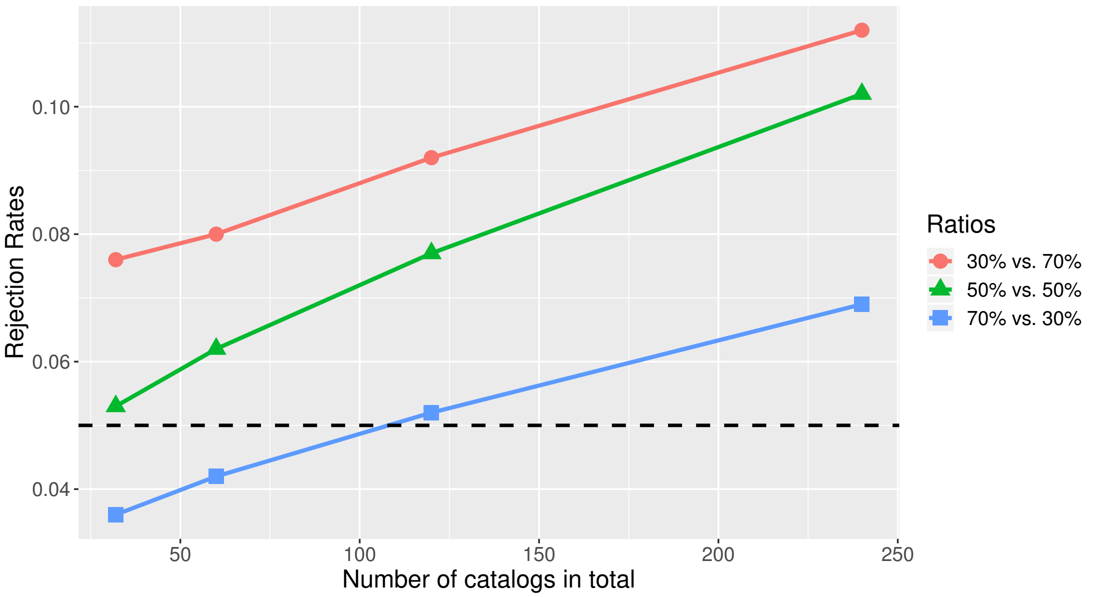


A small simulation study demonstrates how the rejection rate of the two-stage method could be influenced by different variances in two groups with the same means. The rejection rates are defined as the fraction of significant results with Wilcoxon P value less than 0.05 given no mean difference in two groups. To determine the rejection rates, two groups of data were randomly generated from two distributions with the same mean but different variances, Dirichlet(1, 1, 1) and Dirichlet(2, 2, 2). In addition, to better understand how rates are influenced by the number of total catalogs and the sample size ratio between groups, this comparison is repeated after varying the total number of catalogs (32, 60, 120, 240) and sample size ratios between the two groups (30% vs. 70%, 50% vs. 50%, 70% vs. 30%). The simulations were repeated 5,000 times for each scenario.

As shown in the figure, the rates of significant results are inflated by the different variances and worsen along with the increasing number of total catalogs. For any given number of total catalogs, the larger the fraction of catalogs from the distribution with larger variance, e.g., Dirichlet(1, 1, 1), the less likely the test will be significant (blue line).
